# GDNF-RET signaling and EGR1 form a positive feedback loop that promotes tamoxifen resistance via cyclin D1

**DOI:** 10.1101/2022.05.31.492660

**Authors:** Brooke A. Marks, Ilissa M. Pipia, Chinatsu Mukai, Sachi Horibata, Edward J. Rice, Charles G. Danko, Scott A. Coonrod

## Abstract

**Background:** Rearranged during transfection (RET) tyrosine kinase signaling has been previously implicated in endocrine resistant breast cancer, however the mechanism by which this signaling cascade promotes resistance is currently not well described. We recently reported that glial-cell derived neurotrophic factor (GDNF)-RET signaling appears to promote a positive feedback loop with the transcription factor early growth response 1 (EGR1). Here we investigate the mechanism behind this feedback loop and test the hypothesis that GDNF-RET signaling forms a regulatory loop with EGR1 to upregulate cyclin D1 (CCND1) transcription, leading to cell cycle progression and tamoxifen resistance.

**Methods:** To gain a better understanding of the GDNF-RET-EGR1 resistance mechanism, we studied the GDNF-EGR1 positive feedback loop and the role of GDNF and EGR1 in endocrine resistance by modulating their transcription levels using CRISPR-dCAS9 in tamoxifen sensitive (TamS) and tamoxifen resistant (TamR) MCF-7 cells. Additionally, we performed kinetic studies using recombinant GDNF (rGDNF) treatment of TamS cells. Statistical significance for qPCR and chromatin immunoprecipitation (ChIP)-qPCR was determined using a student’s t-test.

**Results:** GDNF-RET signaling formed a positive feedback loop with EGR1 and also downregulated estrogen receptor 1 (ESR1) transcription. Upregulation of GDNF and EGR1 promoted tamoxifen resistance in TamS cells and downregulation of GDNF promoted tamoxifen sensitivity in TamR cells. Additionally, we show that rGDNF treatment activated GDNF-RET signaling in TamS cells, leading to recruitment of p-ELK-1 to the EGR1 promoter, upregulation of EGR1 mRNA and protein, binding of EGR1 to the GDNF and CCND1 promoters, increased GDNF protein expression, and subsequent upregulation of CCND1 mRNA levels.

**Conclusion:** Outcomes from these studies support the hypotheses that GDNF-RET signaling forms a positive feedback loop with the transcription factor EGR1, and that GDNF-RET-EGR1 signaling promotes endocrine resistance via signaling to cyclin D1. Inhibition of components of this signaling pathway could lead to therapeutic insights into the treatment of endocrine resistant breast cancer.

## Background

In estrogen receptor α positive (ERα+) breast cancer (BC), ERα signaling becomes overactive leading to uncontrolled proliferation through activation of proliferative factors, such as cyclins, and enhanced cell survival through inhibition of apoptotic factors [1]. Endocrine therapies are a common tool to inhibit specific aspects of this signaling pathway in BC patients, with tamoxifen (TAM) currently being the most common ERα+ BC therapy. However, about 20-30% of tumors from BC patients are either initially resistant to TAM therapy, also known as *de novo* resistance, or acquire TAM resistance during therapeutic treatment, resulting in tumor progression, metastasis, and increased mortality rates [2].

Several distinct processes have been implicated in TAM resistance (reviewed in [3]), with one being the upregulation of “escape pathways”. One well described escape pathway is epidermal growth factor receptor (EGFR)/HER2 signaling [4]–[7], while a less understood pathway implicated in endocrine resistance is the RET signaling cascade [8]. RET signaling is activated by several ligands, including GDNF. Previous studies have suggested that GDNF/RET tyrosine kinase activates mitogen-activated protein kinase (MAPK) signaling components leading to endocrine resistant ERα+ breast cancer [1], [9]–[11] by promoting cell survival and proliferation that persists in the presence of endocrine therapies. While a relationship between GDNF-RET signaling and endocrine therapy resistance has been previously described [8], [9], [11], the mechanism remains unclear.

At the mechanistic level, GDNF is believed to activate the classical MAPK signaling pathway [12], [13], through association with the RET receptor, leading to extracellular signal-regulated kinases 1 and 2 (ERK 1/2) translocation to the nucleus and phosphorylation of downstream transcription factors (TF), such as Ets Like-1 (ELK-1). Interestingly, ELK-1 is known to bind to the EGR1 promoter, which we have previously shown to be upregulated by GDNF [9]. EGR1 is a transcription factor and member of the immediate-early gene family that is activated through multiple stimuli, including growth factors. Additionally, EGR1 is important for the regulation of cell growth, differentiation, and apoptosis and is known to be a downstream target of MAPK signaling [14]. EGR1 has also been identified as an important component of cancer pathways and can both suppress and promote tumor progression (reviewed in [15]). Further, EGR1 is highly expressed in several different cancers, such as glioma, lung, ovarian, and prostate cancer [15]–[22].

EGR1 has also been linked to endocrine resistance [23], however, the importance of this transcription factor in resistance remains controversial. Importantly, EGR1 has been proposed to be important for cell cycle progression in both MCF-7 TamS and TamR cells [24]. In further support of the role of EGR1 in BC cell proliferation, EGR1 has been shown to directly bind to the CCND1 promoter [25] and the role of the encoded cyclin D1 protein in cell cycle progression and proliferation among multiple cell types is well-described. Interestingly, ERα is also known to promote cell proliferation by binding to CCND1 regulatory regions [26]. Therefore, we hypothesized that upregulation of cyclin D1 through EGR1 may promote cell cycle progression in the presence of endocrine treatment, bypassing the requirement of ERα signaling, to promote resistance.

This current study investigates the relationship between GDNF and EGR1 and how this connection sustains cell proliferation in the presence of tamoxifen. Given the above observations, as well as previous data showing EGR1 upregulates GDNF [9], we hypothesized that GDNF-RET signaling promotes the upregulation of EGR1 expression, and in turn, EGR1 then upregulates GDNF expression, forming a positive feedback loop, with subsequent upregulation of CCND1 expression, thereby promoting cell proliferation and survival in the absence of ERα, leading to endocrine resistance.

To test the hypothesis that GDNF-RET and EGR1 form a positive feedback loop to promote TAM resistance, we initially used CRISPR-dCAS9 to endogenously modulate transcription of GDNF and EGR1. We investigated how transcriptional changes of GDNF altered transcription of EGR1 and vice versa, as well as how these changes altered TAM sensitivity in MCF-7 tamoxifen sensitive (TamS) and *de novo* tamoxifen resistant (TamR) subcloned cell lines (figure 1a). Additionally, we activated RET signaling in TamS cells by treating cells with rGDNF to investigate the mechanism of resistance in a stepwise manner. To test the hypothesis that EGR1 upregulates CCND1 transcription to promote proliferation, we investigated EGR1 binding at the CCND1 promoter after rGDNF treatment. Outcomes from these studies should lead to a deeper understanding of the GDNF-RET signaling mechanism and has the potential to aid in the development of novel therapies, the repurposing of current therapies, and the identification of biomarkers for the treatment of endocrine resistant breast cancer.

**Figure 1.**
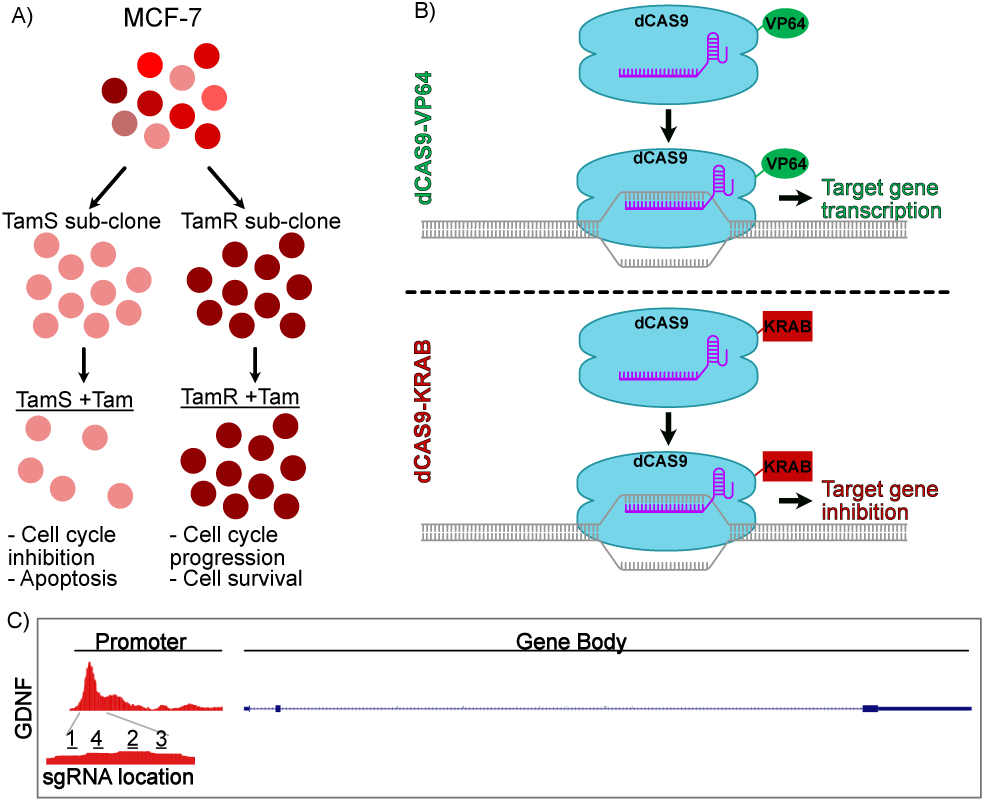
CRISPR-dCAS9 provides the ability to alter gene transcription. (A) Schematic illustration of cell line information. (B) Schematic of dCAS9 system used. (C) Location of sgRNA’s targeting the GDNF promoter.

## Methods

### Cell lines and cell culture

TamS and TamR MCF-7 cells were a gift from Dr. Joshua LaBaer [27]. Cells were grown in Dulbecco’s Modified Eagle Medium (DMEM) with 5% fetal bovine serum (FBS) and 1X Antibiotic-Antimycotic. Reagents used throughout this paper were: Doxycycline hyclate (2ug/ml, Sigma, cat# D9891-5G), Tamoxifen as (Z)-4-Hydroxytamoxifen (4-OHT; 1uM and 5uM, Sigma-Aldrich; Cat# H7904), Recombinant GDNF (10ng/ml, Peprotech, cat# 450-10-10UG). TamS cells were transduced using a lentivirus (Mirus Trans-Lenti Transfection Reagent; Cat # MIR 6600) containing the CRISPR-dCAS9-VP64 (addgene # 50918) for the GDNF upregulated cells and CRISPR-dCAS9-TRE-VP64 (addgene #50916) for the EGR1 upregulated cells. TamR cells were transduced using the same lentivirus, containing the CRISPR-dCAS9-KRAB (addgene # 50919) for the GDNF downregulated cells. The dCAS9-VP64 and dCAS9-KRAB plasmids contained constitutive transcription of the dCAS9-VP64/dCAS9-KRAB genes when stably inserted into the genome. The dCAS9-TRE-VP64 plasmid contained a tetracycline inducible system, where transcription of these genes were induced with doxycycline (dox). The sgRNA plasmids were stably inserted into the genome using the above mentioned lentivirus system. The sgRNA plasmid used for insertion of all sgRNAs was pLenti SpBsmBI sgRNA Hygro plasmid (addgene #62205) and sgRNA sequences are shown in Table S1. All sgRNAs were constitutively expressed. Cells were selected for successful insertion using the selection marker indicated on addgene plasmid information.

### RNA extraction and quantitative real-time PCR (qPCR)

Cells were seeded at 500,000 cells/well in a 6 well plate. RNA was collected from GDNF modulated cells 24hrs later. EGR1-modulated cells were treated with +/- dox for 24hrs prior to RNA collection. For all cell lines, TRI Reagent (Molecular Research Center, Inc., cat #TR118) was used to extract RNA following the manufacturer’s instructions. cDNA was made using High-Capacity RNA-to-cDNA kit (Applied Biosystems, cat #4387406), in accordance with the manufacturer’s instructions and then was diluted 1:5 in molecular grade water. PowerUp SYBR Green was used according to the manufacturers instructions (Life Technologies Cat#A25741). ΔΔCt method was used for all qPCR analyses. Data are represented as mean ±SEM (n≥3) unless otherwise mentioned. qPCR primers are found in Table S1.

### Crystal violet cell proliferation assay

Cells were seeded at 20,000 or 50,000 cells per well on a 12-well plate for all experimentation. GDNF-modulated cell lines were treated with 1 *µ*M TAM, 5 *µ*M TAM, or 100% EtOH vehicle control. Cells were grown for 6 days with or without TAM. EGR1-modulated cell lines were treated with dox or 70% EtOH for the vehicle control and 1 *µ*M TAM. Media was changed every 48hrs. Each cell line was thus treated with +Dox +Tam, +Dox -Tam, -Dox +Tam, and -Dox-Tam. All experiments were performed in triplicate. After 6 days, wells were washed with 1x phosphate-buffered solution (1x PBS), fixed with 500 uL of 2% PFA for 10 minutes, washed with 1x PBS, stained with crystal violet (CV) stain for 10 minutes, and then washed three times with 1x PBS before air drying. Once dry, 500 uL of 10% acetic acid was added to each well. The absorbance of CV in each well was detected by a TECAN plate reader (infinite M200PRO) at 595 nm. Data are represented as mean ±SEM (n≥3) unless otherwise mentioned.

### Protein analysis

Cells were seeded at 500,000 cells per plate, treated with rGDNF, and collected at each specific time point. Cells were washed with ice-cold 1x PBS before lysing with radioimmunoprecipitation assay buffer (RIPA) buffer. Pierce protease inhibitor (Thermo Scientific, ref# A32955) and 10 mM sodium fluoride (NaF), a phosphatase inhibitor, was added to the RIPA buffer. Cells were removed using a cell lifter over ice and spun down at 13,000g at 4°C for 20 minutes. Supernatant was stored at -20°C. A micro bicinchoninic acid (BCA) assay (Thermo Scientific, cat # 23235) was used according to the manufacturer’s instructions to determine protein concentration. The Bio-Rad Trans-Blot Turbo western blot semi-dry transfer system was used for all transfers. Protein was transferred to 0.2um PVDF membrane (IMMUNOBILON-P^SQ^ cat # ISEQ00010). All antibodies were diluted in blocking solution and incubated overnight at 4°C. Anti-EGR1 (B-6)x (santa cruz; cat #sc-515830X) was used at 1:1000 in 5% bovine serum albumin (BSA). GDNF (AbCam, ab18956) and B-Actin (AbCam, ab8227) antibodies were used at 1:125 and 1:5000 in 5% non-fat milk, respectively. Membranes were washed in tris buffered saline with tween (TBST). Secondary anti-rabbit and anti-mouse peroxidase-conjugated affiniPure antibodies from Jackson ImmunoResearch Laboratories (cat #111-035-144 and cat# 115-035-146, respectively) were diluted to 1:20000 and 1:10000, respectively, in TBST and incubated at room temperature for 1hr. Membranes were washed 3x for 20 minutes with TSBT and then imaged using WesternBright Quantum detection kit (cat# K-12042-D10) and Bio-Rad ChemiDoc MP.

### Chromatin Immunoprecipitation

Cells were plated at 5,000,000 cells in 150mm dishes and grown to 90-95% confluence. Cells were treated with +/- 10 ng/ml rGDNF for a specific time. Cells were then washed with PBS, cross-liked with 0.75% paraformaldehyde (Electron Microscopy Sciences, cat#15710) at room temperature for 10 minutes, and the crosslinking was quenched using glycine (125mM final concentration, Fisher Scientific; cat# BP381-5). Cells were then washed twice with ice-cold PBS, harvested in 5ml of PBS with protease and phosphatase inhibitors, spun at 1000g for 5min at 4°C, and the supernatant was discarded. Pellet was resuspended in 325ul ChIP lysis buffer, incubated on ice for 10 minutes, and sonicated for 30 sec on/30 sec off until fragments were between 100-600bp. 100 ul of sample was diluted in 900 ul of dilution buffer (0.05% Tween TBS + fresh Protease inhibitor), beads were added to sample for 1 hr at 4C to minimize non-specific background, then beads were removed, corresponding antibodies were added to each diluted aliquot and incubated overnight at 4C on a nutator. Anti-p-ELK-1 antibody (B4) (Santa Cruz, cat# sc-8406x) was used at 5ug. Anti-EGR1 antibody (B6) (Santa Cruz, cat# sc-515830x) was used at 10ug. Mouse IgG control antibody was from Diagenode (cat# C15400001). Input was kept on a nutator overnight without antibody. Immobilized Protein G (cat# 786-284) and Protein A (cat# 786-283) resin from G-Biosciences were used at a 70:30 ratio. 40ul of slurry was added to samples and incubated for 2hrs at 4C on nutator. Beads were spun down, washed 2x 10 minutes at 4C on nutator with wash buffer (0.1% SDS, 1% Triton X-100, 2mM EDTA pH 8, 20mM Tris-HCL pH 8, and 150mM NaCl) and once with high salt wash buffer (0.1% SDS, 1% Triton X-100, 2mM EDTA pH 8, 20mM Tris-HCL pH 8, and 500mM NaCl). Samples were then eluted in 100ul of fresh elution buffer (1% SDS, 100mM NaHCO3, in molecular grade water) and placed at 65°C at 1200rpm for overnight in Eppendorf mastercycler thermomixer. Samples were then de-crosslinked and RNA was degraded using 2 ul of 10mg/ml Ambion RNAse (Invitrogen, cat# AM2270) for 2 hrs. 10 ul of proteinase K was added and incubated at 65C for overnight to cleave and digest proteins in each sample. Samples were then purified using Omega Bio-Tek E.Z.N.A. Cycle Pure Kit for PCR purification (cat# D6492-02) and eluted in 50ul MQ H2O. qPCR was performed using PowerUp SYBR (applied biosystems, cat# A25742). 2ul of each immunoprecipitated sample was used and input was diluted 1:100 before use. Results were analyzed using fold enrichment.

### Statistics

Student’s Paired T-Test were used in all statistical methods. Samples were normalized to controls prior to analysis. Three independent replicates were used, unless otherwise specified.

## Results

### GDNF modulation alters tamoxifen sensitivity in TamS and TamR MCF-7 cell lines

To further investigate the importance of GDNF in TAM resistance, we used a CRISPR dead CAS9 (dCAS9) construct (figure 1b) that targets the GDNF promoter to endogenously upregulate and downregulate GDNF in TamS and TamR cells, respectively. The promoter region (shown in figure 1c) was identified using dREG-HD [28], which shows divergent transcription, a hallmark of promoter and enhancer regions. In TamS cells, dCAS9-VP64 was stably integrated into the DNA, where it was utilized to decondense chromatin and upregulate target gene transcription (figure 1b). Likewise, a dCAS9-KRAB vector was stably incorporated into the TamR cell line, where it functioned to condense chromatin and inhibit target gene transcription (figure 1b). The dCAS9-sgRNA complex was targeted to the GDNF promoter using four constitutively transcribed sgRNA sequences (figure 1c).

TamS cells containing the dCAS9-VP64 and GDNF sgRNAs (TamS^GDNF-OE^) endogenously upregulated GDNF when compared to the dCAS9-VP64 sgRNA vector control (TamS^Vc^) (fig. 2a; p < 0.03). Subsequent crystal violet (CV) cell proliferation assays showed TamS^GDNF-OE^ cells proliferated and/or survived 1.5 times more than TamS^Vc^ cells when treated with 1uM (p = 0.05) and 5uM (p = 0.05) TAM (figure 2e).

**Figure 2.**
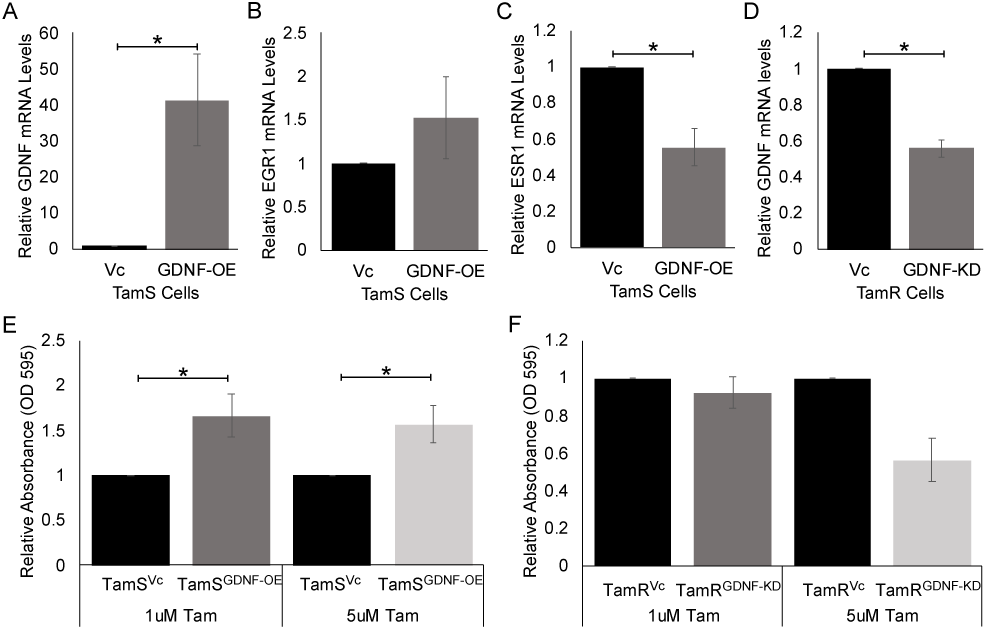
GDNF modulation using CRISPR dCAS9 alters transcription and TAM sensitivity in TamS and TamR cells. (A) GDNF; p<0.05, (B) EGR1; not significant, and (C) ESR1; p<0.006, mRNA expression after endogenous upregulation of GDNF using CRISPR-dCAS9-VP64 system. (D) GDNF mRNA expression after endogenous downregulation of GDNF using CRISPR-dCAS9-KRAB system. (E) Cell Viability of TamS^GDNF-OE^ and TamS^Vc^ cells in the presence of 1uM and 5uM TAM, P<0.05. (F) Cell Viability of TamR^GDNF-KD^ and TamR^Vc^ cells in the presence of 1uM TAM and 5uM (p < 0.065) TAM. Data in (F) are represented as mean ±SEM (n=2). Student’s paired t-test was used for statistical analyses.

Moreover, TamR cells containing the dCAS9-KRAB and GDNF sgRNAs (TamR^GDNF-KD^) endogenously downregulated GDNF transcription when compared to the dCAS9-KRAB sgRNA vector control (TamR^Vc^) (fig 2d, p < 0.001). Downregulation of GDNF in TamR^GDNF-KD^ cells promoted tamoxifen sensitivity compared to the TamR^Vc^ after 5uM of TAM treatment (p < 0.065), but not after 1uM of TAM treatment (fig. 2f).

### Endogenous upregulation of GDNF and EGR1 alters transcription

We found that sustained upregulation of GDNF in TamS^GDNF-OE^ cells upregulated EGR1 transcription by 2-fold and downregulated ESR1 by 0.4 fold (p < 0.005) (fig. 2b and 2c). In order to further investigate the relationship between EGR1 and GDNF signaling, we upregulated EGR1 in TamS cells (TamS^EGR1-OE^). Due to the variable nature of EGR1 expression [29] and the potential difficulty to achieve sustained upregulation, we utilized a tetracycline inducible dCAS9-VP64 system to better upregulate EGR1 expression. In this system dCAS9 expression is dependent upon doxycycline (dox) treatment. Six sgRNAs targeted the EGR1 promoter (fig. 3a). Treatment of TamS^EGR1-OE^ and TamS^Vc^ cells, with and without dox for 24hrs, successfully upregulated EGR1 transcription by 1.5-fold (p < 0.005) and GDNF by 2.5-fold (p < 0.05) (fig. 3c and 3d). Together this data suggests that GDNF and EGR1 form a positive feedback loop (figure 3b). Cell proliferation assays demonstrated that TamS^EGR1-OE^ cells proliferated 2 times faster than TamS^Vc^ cells in the presence of 1uM TAM, however this result was not statistically significant, potentially due to variability within replicates (Figure S1).

**Figure 3.**
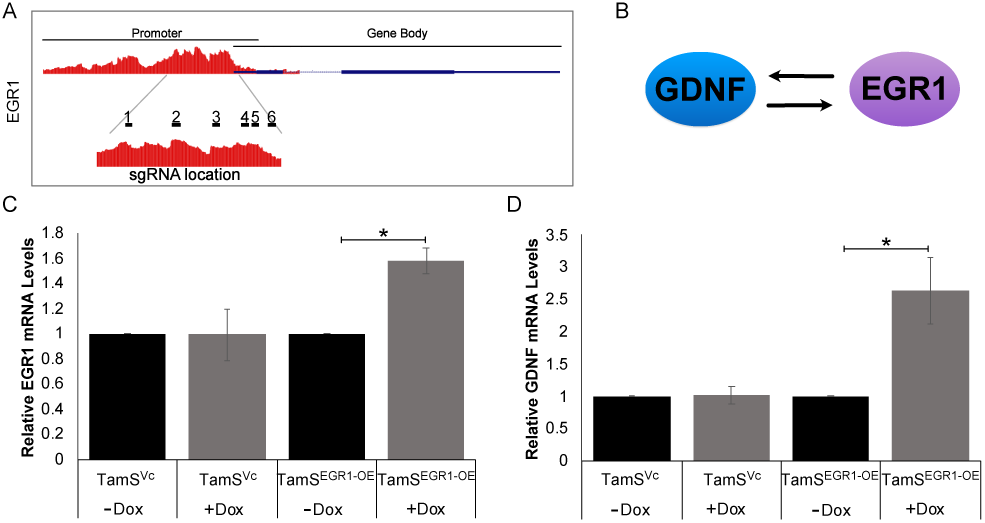
TamS^EGR1-OE^ cells upregulates GDNF transcription. (A) Location of sgRNA’s targeting the EGR1 promoter. (B) Schematic of GDNF-EGR1 feedback loop. (C) EGR1 (p<0.005) and (D) GDNF (p<0.05) mRNA expression levels after 24hrs of dox treatment in TamS^EGR1-OE^.

### Recombinant GDNF (rGDNF) treatment activates the GDNF-EGR1 positive feedback loop in TamS cells

TamS cells express RET tyrosine kinase but do not secrete the GDNF ligand [10]. Addition of the RET ligand rGDNF activated the RET signaling pathway *in vitro*, thereby allowing us to test the effect of GDNF treatment on EGR1 expression over time. Results showed that rGDNF treatment significantly upregulated EGR1 mRNA expression in TamS cells at 2, 3, 6, and 12 h post treatment (figure 4a). Additionally, we found that treatment of TamS cells with rGDNF also induced the expression of EGR1 protein at 1h, with levels of EGR1 protein further increasing at subsequent timepoints. rGDNF treatment also induced endogenous GDNF expression beginning at ∼3h post treatment, with protein levels appearing to increase at subsequent hours (figure 4b). Interestingly, higher MW protein bands were stained with the EGR1 antibody, suggesting that rGDNF treatment may also promote EGR1 phosphorylation (potentially resulting in EGR1 activation).

**Figure 4.**
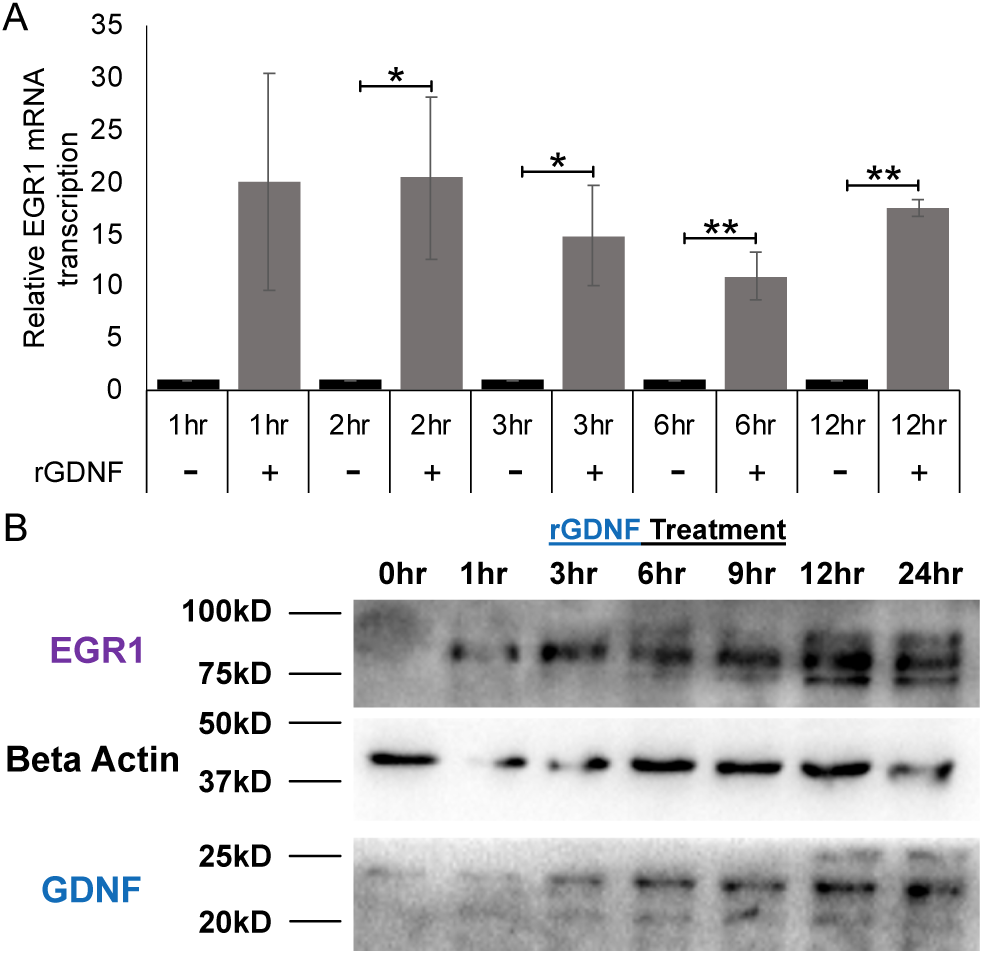
rGDNF upregulates EGR1 and GDNF. (A) EGR1 mRNA expression levels after rGDNF treatment. Student’s paired t-test was used for statistical analyses. * p < 0.05, ** p < 0.005. (B) Cropped blots of EGR1, GDNF, and B-Actin protein expression after rGDNF treatment at various treatment times. Uncropped blots are presented in Figure S2.

### GDNF upregulates EGR1 transcription through ELK1 phosphorylation

GDNF-RET signaling has been shown to activate the MAPK signaling cascade, with ELK-1 and the transcription factor SRF (serum response factor) both being downstream targets of this kinase [9], [30]. Interestingly, through analysis of existing datasets using the UCSC genome browser, we identified an SRF ChIP-seq peak and ELK-1 binding motifs located at the EGR1 promoter in MCF-7 cells (fig. 5a). To further understand the mechanism by which EGR1 is upregulated, we performed a Chromatin Immunoprecipitation (ChIP)-qPCR assay using an anti-phospho-ELK-1 antibody to investigate ELK-1 phosphorylation and binding at the EGR1 promoter following GDNF treatment. Results showed that treatment of TamS cells with rGDNF resulted in a four fold enrichment of p-ELK-1 (p < 0.005) at the EGR1 promoter compared to untreated cells (fig. 5b). This result suggests that GDNF-RET signaling promotes ELK-1 binding and phosphorylation at the EGR1 promoter.

**Figure 5.**
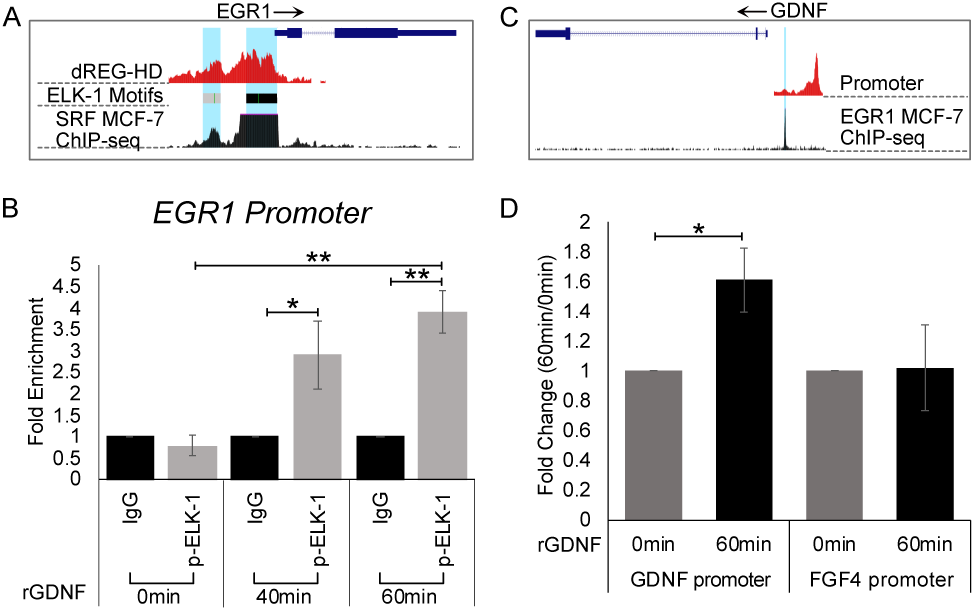
Mechanism of GDNF-EGR1 positive feedback loop. (A) UCSC genome browser data showing p-ELK-1 binding motif located at EGR1 promoter. (B) Fold enrichment of ChIP-qPCR of anti-p-ELK-1 antibody binding at the EGR1 promoter. (C) UCSC genome browser data showing EGR1 binding motif located at the GDNF promoter. (D) ChIP-qPCR of anti-EGR1 anitbody targeting the GDNF promoter. *p < 0.05, **p < 0.005. Student’s paired t-test was used for statistical analyses.

### EGR1 directly binds to the GDNF promoter

Analysis of UCSC genome browser data identified an EGR1 ChIP-seq peak at the GDNF promoter in MCF-7 cells (fig. 5c). This data, with the above results, suggests that rGDNF treatment may promote EGR1 binding at the GDNF promoter. We tested this hypothesis and found that treatment with rGDNF resulted in a 0.6 fold enrichment of EGR1 binding to the GDNF promoter after 60 minutes of rGDNF treatment when compared to the 0hr control (figure 5d, p < 0.05). This enrichment was not observed in the negative control FGF4 promoter region that does not contain and EGR1 binding motif.

### The GDNF-EGR1 feedback loop likely promotes endocrine resistance by inducing CCND1 transcription

Our study suggests that GDNF and EGR1 appear to form a positive feedback loop, however, how this feedback loop promotes TAM resistance is unknown. Analysis of UCSC genome browser data from MCF-7 cells identified an EGR1 ChIP-seq peak and four EGR1 binding motifs at the CCND1 promoter (figure 6a). Moreover, a previous study [25] suggests that EGR1 can bind to the CCND1 promoter to induce transcription. Therefore, we hypothesized that the GDNF-EGR1 positive feedback loop sustains EGR1 expression, and in doing so, EGR1 binds to the CCND1 promoter to induce transcription and promote cell proliferation in the presence of TAM. To test if EGR1 binds to the CCND1 promoter, we performed two experiments. First, TamS cells were treated with TAM for 24hrs to inhibit cell proliferation and CCND1 transcription. Cells were then treated for various times with or without rGDNF, still in the presence of TAM. After 2, 3, and 6 h of rGDNF treatment, CCND1 transcription was significantly upregulated compared to the control (figure 6b, p < 0.05). Additionally, ChIP-qPCR was performed using an anti-EGR1 antibody after 60 minutes of rGDNF treatment resulting in increased EGR1 binding to the CCND1 promoter by 1.6 fold (figure 6c, p < 0.05). This enrichment was not observed when compared to the FGF4 promoter control.

**Figure 6.**
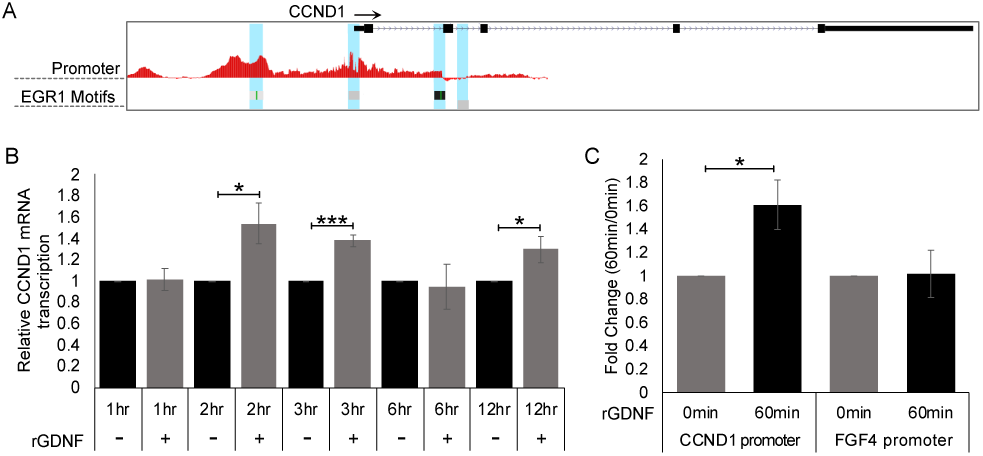
CCND1 upregulation through GDNF-RET signaling. (A) UCSC genome browser data showing EGR1 binding motif located on the CCND1 promoter. (B) CCND1 mRNA levels after 1uM TAM treatment and +/- rGDNF treatment. (C) ChIP-qPCR of anti-EGR1 antibody targeting the CCND1 promoter. Student’s paired t-test was used for statistical analyses. * p< 0.05, *** p < 0.0005.

## Discussion

The development of therapeutic resistance in cancer leads to progression, metastasis, and decreased overall survival. Understanding the mechanism of action behind resistance is critical in developing and repurposing current therapies. In ERα+ BC, there are multiple mechanisms of resistance, some of which remain unclear and/or unknown. Among these mechanisms, RET signaling has been implicated in other cancers as well as breast cancer resistance (reviewed in [31]), however, the mechanism behind how RET signaling promotes resistance in ERα+ BC has not been previously studied. In this current study, we used TamS and TamR MCF-7 cells to elucidate the GDNF-RET signaling resistance mechanism of action.

Initially, we found that endogenous modulation of GDNF using the CRISPR-dCAS9 system altered tamoxifen sensitivity in ERα+ MCF-7 subclones. More specifically, TamR^GDNF-KD^ cells behaved similar to TamS cells and TamS^GDNF-OE^ cells behaved similar to TamR cells. This switch in tamoxifen sensitivity has been observed in other studies using BC cell lines, where inhibition of endocrine resistant mechanisms, like GDNF-RET-EGR1 signaling, re-sensitizes cells to TAM. For example, inhibition of c-Cbl, c-src, and HER2 in BT474 cells [32] and inhibition of RET and EGFR (using Gefitinib) in MCF-7 cells [8] [33], re-sensitized cells to TAM. This suggests that there is crosstalk between ERα signaling and mechanisms of resistance, such as GDNF-RET signaling.

Moreover, we found that in TamS^GDNF-OE^ cells, EGR1 transcription was upregulated and ESR1 transcription was downregulated, suggesting that GDNF signaling promotes endocrine resistance through simultaneously downregulating ESR1 and upregulating EGR1, thereby prompting a switch in the signaling pathway used for cell proliferation and survival. Further investigation of this crosstalk and studies using TAM with other therapies to inhibit TAM resistance from occurring and/or resensitize tumors to TAM is needed. We also found that TamS^EGR1-OE^ cells upregulated GDNF transcription and were more resistant to TAM, though this was not significant. Taken together, the above data suggests that EGR1 upregulates GDNF transcription, and GDNF-RET signaling upregulates EGR1 transcription, supporting the idea that GDNF and EGR1 positively regulate each other.

To further investigate the GDNF-RET signaling positive feedback loop mechanism, we performed kinetic studies in TamS cells. We observed that in the presence of both TAM and rGDNF treatment, EGR1 transcription was upregulated and sustained upregulation throughout treatment. Also, in further support of the GDNF-EGR1 positive feedback loop, EGR1 and GDNF protein expression was observed following rGDNF treatment and protein expression then continued to increase throughout the rGDNF treatment period.

We next investigated how EGR1 was upregulated through GDNF-RET signaling. In the MAPK signaling cascade, it is known that ERK1/2 translocates to the nucleus, where it phosphorylates multiple targets, including ELK-1. Our results show that rGDNF treatment led to increased phosphorylation of ELK-1 bound to the EGR1 promoter. These findings fit well with a previous study showing EGR1 activation through the GDNF-RET-MAPK signaling cascade [9].

As previously mentioned, upregulation of GDNF protein was observed following upregulation of EGR1 protein expression. Interestingly, UCSC genome browser data shows an EGR1 binding motif and EGR1 MCF-7 ChIP-seq binding located at the GDNF promoter. Using ChIP-qPCR, we determined that EGR1 binds directly to the GDNF promoter following rGDNF treatment, completing the GDNF-RET-EGR1 positive feedback loop. The formation of a positive feedback loop with EGR1 has also been previously identified in non-cancerous cell activity where EGR1 forms a positive feedback loop with osteopontin (OPN)-MAPK signaling through direct binding of EGR1 to the OPN promoter in vascular smooth muscle cells [34]. Our findings show that in TamR MCF-7 cells, the GDNF-RET-EGR1 positive feedback loop has likely been exploited to promote TAM resistance in TamR MCF-7 cells.

We predict that this positive feedback loop represents one mechanism used in the progression of endocrine resistant BC. This prediction is supported by the observation that RET expression is observed in 60% in patients with recurrent disease after receiving adjuvant tamoxifen therapy [8]. Further investigation into the expression of GDNF and EGR1 in RET positive BC tissue samples is needed to determine the clinical relevance of this potential resistance mechanism.

We predict that inhibition of RET signaling could represent a new therapeutic option for the treatment of endocrine resistant BC, as inhibition of RET signaling through GDNF inhibition promoted TAM sensitivity. Interestingly, the first RET inhibitor selpercatinib (Retevmo) was recently approved to treat non-small-cell lung cancer and two forms of thyroid cancer that contain RET alterations [35]. Further investigations are warranted to determine if ERα+ BC patients harbor these alterations, and if Retevmo could be a potential therapeutic, either alone or with endocrine therapies.

Regarding our findings with CCND1, we predicted, that upon ERα inhibition, GDNF-RET signaling upregulates EGR1, not only to form a positive feedback loop with GDNF, but also to promote cell proliferation in ERα+ BC patients through EGR1 binding directly to the CCND1 promoter to induce CCND1 transcription. Our results show that in TamS cells, EGR1 not only upregulates GDNF, but also upregulates CCND1 transcription by directly binding to the CCND1 promoter after rGDNF treatment. Additionally, in TamS cells treated with TAM, CCND1 transcription was upregulated by the presence of rGDNF. This data suggests that GDNF-RET signaling promotes cell proliferation through upregulation of the transcription factor EGR1, which in turn upregulates both GDNF to promote a positive feedback loop and CCND1 to ultimately promote cell cycle progression and survival (figure 7). Given our findings, one potential therapeutic option to treat GDNF-RET-EGR1 resistant BC cancer would be to inhibit cyclin D1-CDK 4/6. Venzenio, also known as abemaciclib, is a CDK 4/6 inhibitor that has been approved by the FDA to treat advanced or metastatic breast cancers [36]. This drug is used for the treatment of hormone receptor positive, HER2 negative BC. Further investigation is needed into Venzenio as a potential therapeutic treatment for endocrine resistant BC that has activated GDNF-RET signaling.

**Figure 7.**
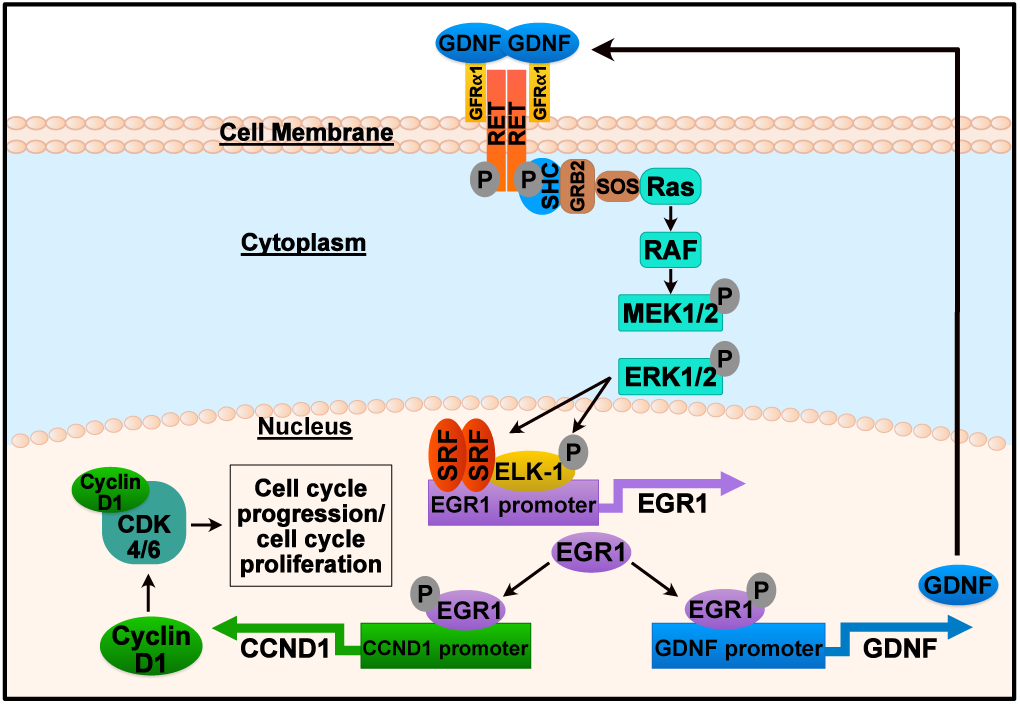
Proposed mechanism for TAM resistance in ERα+ breast cancer. (A) Graphical summary of one tamoxifen resistant mechanism in ERα+ breast cancer where GDNF-RET-EGR1 signaling form a positive feedback loop and promote cell cycle progression through CCND1 upregulation.

## Conclusion

Overall, we observed that GDNF-RET signaling forms a positive feedback loop with EGR1, and in turn, EGR1 upregulates CCND1 to induce cell proliferation, therefore promoting tamoxifen resistance. Inhibition of this signaling pathway, either through RET, MAPK, or CCND1 inhibition, could have a positive impact on treating advanced endocrine resistant BC patients. Further studies are needed to determine the prognostic impact of these inhibitors in GDNF-RET-EGR1 expressing BC and whether these inhibitors should be used alone or in combination with current endocrine therapies.

## Declarations

### Ethics approval

Not applicable.

### Consent for publication

The authors consent to publication of this manuscript in BMC Cancer.

### Availability of data and materials

Data files for dREG analysis have been deposited in Gene Expression Omnibus under Accession Number GSE93229. ChIP-seq data was retrieved from publicly available MCF-7 data from the ENCODE project.

### Author Contributions

B.A.M., C.G.D, and S.A.C. participated in the conceptualization, design, and manuscript revisions. B.A.M., C.M, C.G.D, and S.A.C. contributed to data interpretation. B.A.M, I.M.P, S.H., and E.J.R. contributed to methodology and analysis. B.A.M. wrote the main manuscript text and prepared figures. All authors reviewed the manuscript.

## Acknowledgements

The authors thank Marlena Holter for performing preliminary experiments that led to experiments published in this paper and Kelly Sams for providing editing expertise in the draft of this manuscript.

## Abbreviations

BC: Breast cancer
ERα: estrogen receptor alpha
GDNF: Glial cell-derived neurotrophic factor
RET: rearranged during transfection
EGR1: early growth response 1
MAPK: mitogen-activated protein kinase
ERK 1/2: extracellular signal-regulated kinases 1 and 2
CRISPR: Clustered Regularly Interspaced Short Palindromic Repeats
dCAS9: Endonuclear deficient CRISPR-associated protein 9
ELK-1: Ets Like-1
TF: Transcription factor
TAM: Tamoxifen
TamS: Tamoxifen sensitive
TamR: Tamoxifen resistant
HER2: human epidermal growth factor receptor 2
EGFR: Epidermal growth factor receptor

