## Supplemental Figures for "GDNF-RET signaling and EGR1 form a positive feedback loop that promotes tamoxifen resistance via cyclin D1"

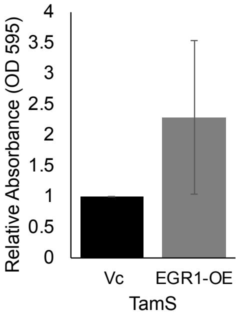


Supplemental Figure 1. Cell Viability of $TamS^{EGR1\text{-}OE}$ and $TamS^{Vc}$ cells in the presence of 1uM TAM. Data in are represented as mean ±SEM.

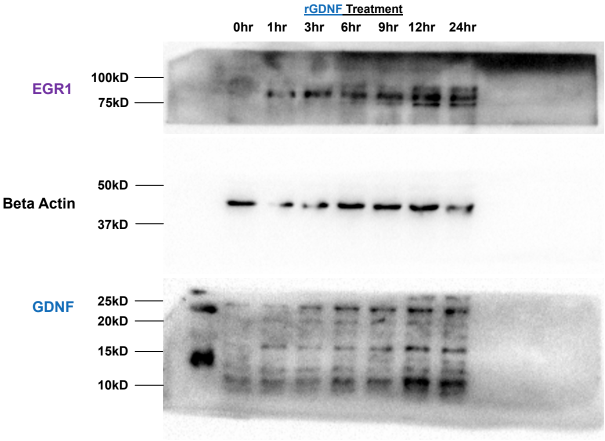


Supplemental Figure 2. Uncropped western blot from figure 4.
