## Supplemental Table 1 for "GDNF-RET signaling and EGR1 form a positive feedback loop that promotes tamoxifen resistance via cyclin D1"

**Table S1. sgRNA and primer sequences**

| Primer | Sequence |
|---|---|
| GDNF sgRNA 1F | acacCGTGCCAGGGGACGCTGCGAGTG |
| GDNF sgRNA 1R | aaaaCACTCGCAGCGTCCCCTGGCACG |
| GDNF sgRNA 2F | acacCGGACCGTGTTGGCCTTAGCATG |
| GDNF sgRNA 2R | aaaaCATGCTAAGGCCAACACGGTCCG |
| GDNF sgRNA 3F | acacCGCATATCATGTATTACATGGCG |
| GDNF sgRNA 3R | aaaaCGCCATGTAATACATGATATGCG |
| GDNF sgRNA 4F | acacCGAGCCACTGGAGGGCACGTCAG |
| GDNF sgRNA 4R | aaaaCTGACGTGCCCTCCAGTGGCTCG |
| EGR1 sgRNA 1F | acacCGGGATGACAGCGATAGAACCCG |
| EGR1 sgRNA 1R | aaaaCGGGTTCTATCGCTGTCATCCCG |
| EGR1 sgRNA 2F | acacCGCGACCCGGAAATGCCATATAG |
| EGR1 sgRNA 2R | aaaaC TATATGGCATTTCCGGGTCGCG |
| EGR1 sgRNA 3F | acacCGCCTAGAGCTCTAGCCGCCGCG |
| EGR1 sgRNA 3R | aaaaCGCGGCGGCTAGAGCTCTAGGCG |
| EGR1 sgRNA 4F | acacCGCGGCCGGTCCTGCCATATTAG |
| EGR1 sgRNA 4R | aaaaCTAATATGGCAGGACCGGCCGCG |
| EGR1 sgRNA 5F | acacCGCGCCTCCGTCGTGACGTACAG |
| EGR1 sgRNA 5R | aaaaCTGTACGTCACGACGGAGGCGCG |
| EGR1 sgRNA 6F | acacCGGATCCCAGCGCGCAGAACTTG |
| EGR1 sgRNA 6R | aaaaCAAGTTCTGCGCGCTGGGATCCG |
| GDNF mRNA (boulay) F | TCTGGGCTATGAAACCAAGGA |
| GDNF mRNA (boulay) R | GTCTCAGCTGCATCGCAAGA |
| EGR1 mRNA F | GACCGCAGAGTCTTTTCCTG |
| EGR1 mRNA R | AGCGGCCAGTATACGTGATG |
| CCND1 mRNA F | CCCTCGGTGTCCTACTTCAA |
| CCND1 mRNA R | CTCCTCGCACTTCTGTTCCT |
| B-actin mRNA F | CCAACCGCGAGAAGATGA |
| B-actin mRNA R | CCAGAGGCGTACAGGGATAG |
| GDNF promoter F | CACAGAAGTGCTCGCAGAAG |
| GDNF promoter R | GTGGTGGTTCTCCGGTTTTA |
| EGR1 promoter F | GACCCGGAAATGCCATATAA |
| EGR1 promoter R | CTTCTTCCCTCCTCCCAGAG |
| CCND1 promoter F | CACACGGACTACAGGGGAGT |
| CCND1 promoter R | ACTCTGCTGCTCGCTGCTA |
| FGF4 promoter F | TGAAAGGACAGGTAGCAGGG |
| FGF4 promoter R | TGCTAGATTGAGGGCTCTGG |
